# Computational study of the furin cleavage domain of SARS-CoV-2: delta binds strongest of extant variants

**DOI:** 10.1101/2022.01.04.475011

**Authors:** M. Zaki Jawaid, A. Baidya, S. Jakovcevic, J. Lusk, R. Mahboubi-Ardakani, N. Solomon, G. Gonzalez, J. Arsuaga, M. Vazquez, R.L. Davis, D.L. Cox

## Abstract

We demonstrate that AlphaFold and AlphaFold Multimer, implemented within the ColabFold suite, can accurately predict the structures of the furin enzyme with known six residue inhibitory peptides. Noting the similarity of the peptide inhibitors to polybasic furin cleavage domain insertion region of the SARS-CoV-2, which begins at P681, we implement this approach to study the wild type furin cleavage domain for the virus and several mutants. We introduce mutations *in silico* for alpha, omicron, and delta variants, for several sequences which have been rarely observed, for sequences which have not yet been observed, for other coronaviruses (NL63, OC43, HUK1a, HUK1b, MERS, and 229E), and for the H5N1 flu. We show that interfacial hydrogen bonds between the furin cleavage domain and furin are a good measure of binding strength that correlate well with endpoint binding free energy estimates, and conclude that among all candidate viral sequences studied, delta is near the very top binding strength within statistical accuracy. However, the binding strength of several rare sequences match delta within statistical accuracy. We find that the furin S1 pocket is optimized for binding arginine as opposed to lysine. This residue, typically at sequence position five, contains the most hydrogen bonds to the furin, and hydrogen bond count for just this residue shows a strong positive correlation with the overall hydrogen bond count. We demonstrate that the root mean square backbone C-alpha fluctuation of the first residue in the furin cleavage domain has a strong negative correlation with the interfacial hydrogen bond count. We show by considering the variation with the number of basic residues that the maximum mean number of interfacial hydrogen bonds expected is 15.7 at 4 basic residues.

## Introduction

While the spike protein of the SARS-CoV-2 virus is similar to SARS-CoV-1, a key difference is a polybasic insertion beginning at P681 in the spike protein (*1*). It has been shown that this insertion is critical to the higher transmissibility of SARS-CoV-2 (*2*, *3*) over SARS-CoV-1, and that the mutations P681H for the alpha and omicron variants and P681R for the delta variant play a large role in increased transmissibility of the variants over the wild type (WT) (*4*). Similar polybasic furin cleavage domains (FCDs) occur in other human coronaviruses OC43, HUK1, 229E, MERS, and NL63 (*5*), and in many other viruses including H5N1 influenza (*6*).

The FCD of SARS-CoV-2 has not been well studied for at least two reasons. First, the FCD belongs to a rapidly fluctuating random coil region of the protein which has not been resolved by structural probes (see, e.g., Ref. (*7*), PDB structure 7A94, for which residues 677-688 are unresolved). Second, because the furin rapidly cleaves the protein at this domain, there are no bound structures available. The absence of structural data has limited computational studies of the binding domain.

A number of small peptides that can act as furin inhibitors have been studied elsewhere. It is known that the four amino acid inhibitor RVKR, suitably terminated, is a potent inhibitor of furin activity (*8*). Right handed hexa-arginine and nona-arginine peptides are potent inhibitors of furin also (*9*). Additionally, the peptide Arg-Arg-Arg-Val-Arg-4-aminomethyl-benzamidine (RRRVR-Amba, I1 peptide) (*10*), is similar to the delta variant FCD RRRARS, and binds to furin with pM affinity. This leads to the conjecture that the FCDs of SARS-CoV-2 and other coronaviruses may bind in similar fashion to the furin enzyme. For SARS-CoV-2, the insertion that begins with P681 for the WT, alpha, delta, and omicron variants commences a six residue sequence (through 686) that hijacks the furin enzyme from its useful physiological functions to assisting the virus. We have focused on several six amino acid FCDs for SARS-CoV-2 and other viruses.

In the absence of structural data for the FCD, we turned to the deep learning based AlphaFold program (*11*). We used AlphaFold Multimer (*12*), as implemented within the ColabFold environment (*13*), to generate candidate structures for the FCD-furin complexes. We find that AlphaFold accurately predicts the furin structure and the backbone of the bound furin-I1 structure (Figs. 1A,B), so it is natural to attempt binding to the FCD, shown for WT in Fig 1C. We have used AlphaFold Multimer as the only way to generate a *de novo* structure for the WT FCD to furin binding. With this hypothesis, we can either generate de novo structures from AlphaFold Multimer, or assume the WT is well represented by the AlphaFold candidate structure, and mutations from that structure can be used to assess the binding of the FCDs for variants and other viruses. We simulate these structures with molecular dynamics to assess equilibrium binding strength, characterized by two quantities, interfacial hydrogen bonds between the furin and FCD (FCD-furin HBonds), and Generalized Born Surface Area (GBSA) binding energies. Details of the simulation protocols are in the Supplementary Materials and Methods section 0.4. The AlphaFold approach reaches different conclusions about the FCD-furin bound structure than an earlier approach based upon docking (*14*).

**Fig. 1.**
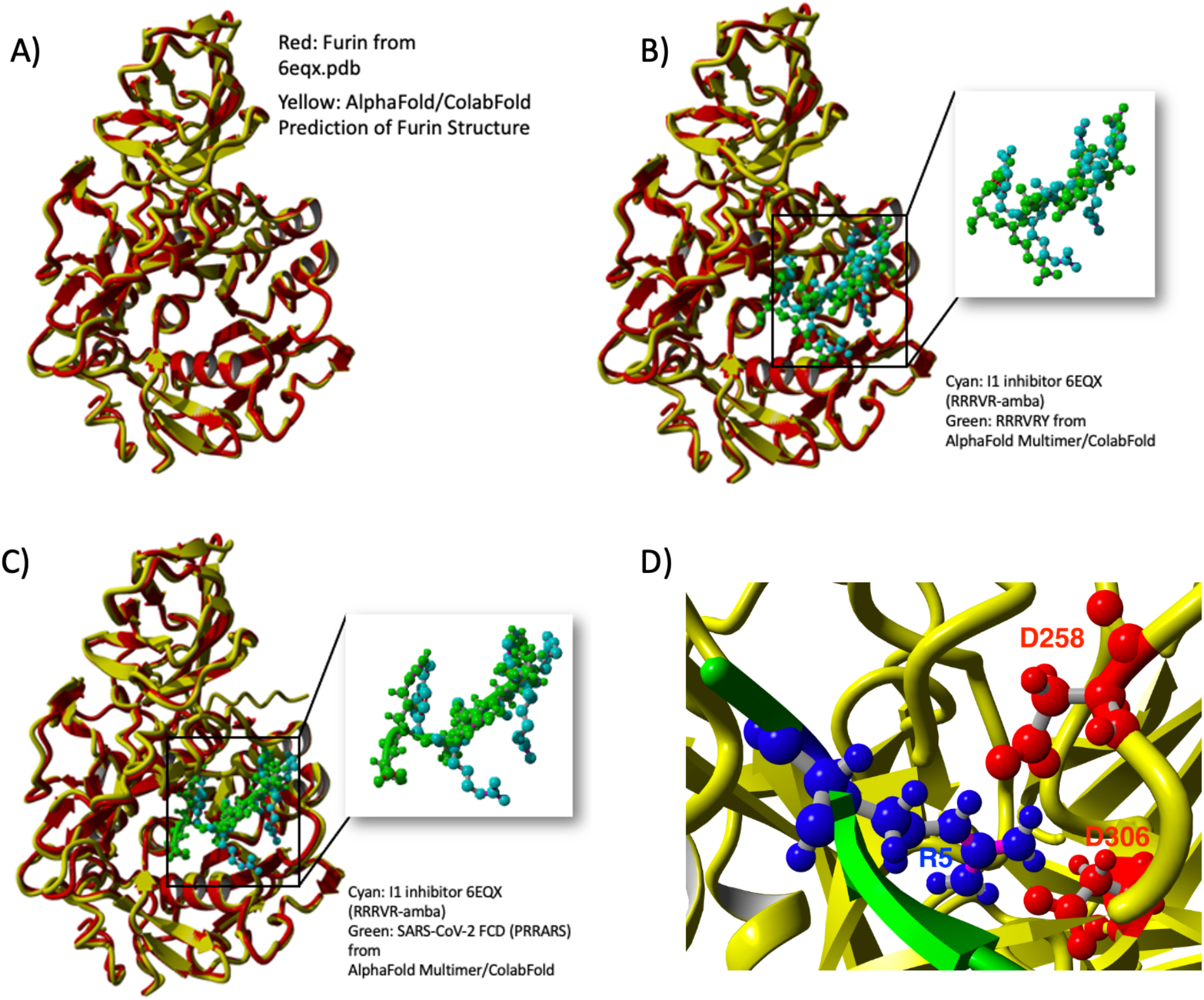
A) Comparison of structure of furin from Ref. (*10*) and PDB file 6EQX with the structure from AlphaFold (*11*) using the ColabFold environment (*13*). Clearly, the agreement is excellent (RMSD of 1.79Å). B) Comparison of structure of furin with RRRVR-Amba inhibitor from Ref. (*10*) with structure generated for the similar sequence RRRVRY by AlphaFold Multimer (*12*) using ColabFold (*13*). The Amba is buried in the furin S1 pocket (*10*) for the inhibitor, while AlphaFold predicts burial of the R at position 5. The backbone RMSD between the I1 and RRRVRY peptides is 2.77Å. C) Predicted structure by AlphaFold Multimer (*12*) for the WT PRRARS sequence of SARS-CoV-2 compared to Furin-I1 structure. D) Close up of binding pocket for fifth residue of PRRARS (WT FCD). Furin backbone in yellow, FCD backbone in green, R5 from FCD is blue, D258,D306 from furin in red.

In the I1 sequence, the sixth (Amba) residue, a nonstandard amino acid, binds most strongly to furin as we discuss below. When we mutate that nonstandard residue to the structurally similar tyrosine, the binding pocket is occupied by the arginine at sequence position 5. Accordingly, we hypothesize that insertion of the residue at position 5 into the furin S1 pocket is the most important for FCD binding to furin. We confirm this hypothesis by simulating dozens of observed sequences. In 93% of observed SARS-CoV-2 FCD sequences starting from aligned position 681, the fifth amino acid is arginine.

We obtain a number of important results. First, per Fig. 2, the delta variant has the strongest binding of existing extant SARS-CoV-2 variants, and only two rare or unobserved FCD sequences bind as strongly within statistical accuracy. This dominance of the delta variant FCD extends to other coronaviruses and the H5N1 influenza virus. Second, as made clear in the heat map of Fig. 3, the most important residue is the fifth, which binds in the S1 pocket of furin (*10*) containing two aspartic acid residues. In particular, this pocket matches structurally to arginine better than lysine as discussed below. Third, we find that there is mechanistic predictive power in three quantities that help explain the differences between delta and other variants and viruses: (*1*) the strength of the binding strongly correlates inversely with the root mean square fluctuation (RMSF) of the first residue. This suggests that the more the backbone outside the pocket fluctuates, the less likely the arginine at position 5 can bind well to the furin. (*2*) The number of FCD-furin hydrogen bonds between residue 5 and the furin strongly predicts the total binding strength, even though it only represents a plurality of the HBonds. (*3*) The maximum mean number of FCD-furin HBonds for a given number of basic residues peaks at 15.7 hydrogen bonds for 4.06 basic residues.

**Fig. 2.**
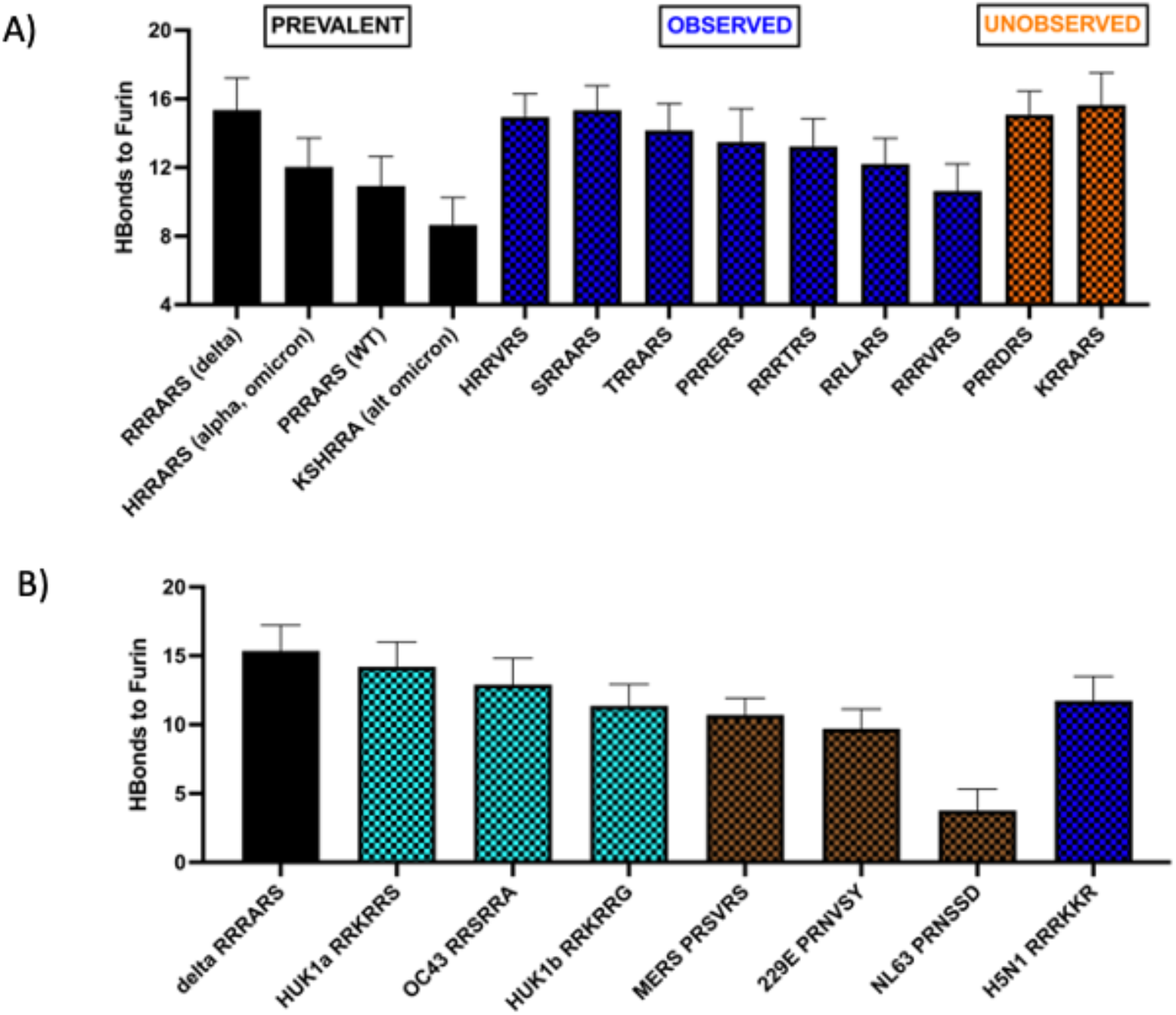
A) FCD-furin hydrogen bond counts between furin and SARS-CoV-2 binding sequences at 681-686 of the spike protein. The first four bars are prevalent forms (WT, delta, omicron/alpha, and alt omicron where we assume the sequence starts at position 679. The blue sequences are rare but observed in the GISAID (*19*) database; of these HRRARN and SR-RARS bind as strongly to furin with in statistical accuracy as the delta sequence (RRRARS). The two unobserved sequences require double base mutations from existing extant codons, but bear watching because of their strong binding to furin. B) FCD-furin hydrogen bond counts between furin and other viruses. The SARS-CoV-2 delta variant shows the strongest binding of any human coronavirus and exceeds the H5N1 influenza cleavage site.

**Fig. 3.**
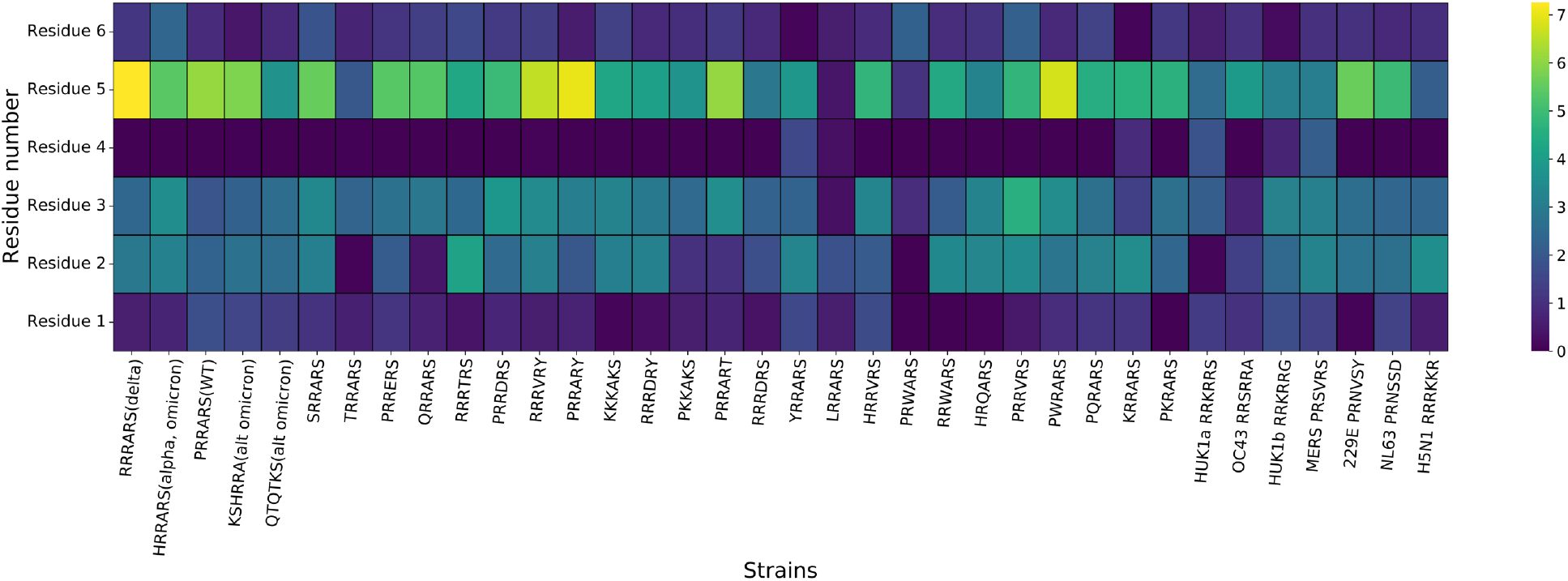
Heat map of interfacial hydrogen bonds from furin to the six residue peptide by residue number (vertical) for various observed SARS-CoV-2 along with two unobserved, and for other human coronaviruses and H5N1. Clearly, the key residue for binding is the fifth.

## Results

To avoid confusion between the conventional N-to-C terminal sequence numbering of peptides and proteins vs. the reverse numbering used in the Furin Data Base (FurinDB) (*15*) and other references (*8*–*10*), we will refer to the FCD residues as positions 1-6, which for all viral sequences considered here will correspond to the FurinDB notation P5-P4-P3-P2-P1-P1’, with the cleavage site between P1 and P1’. For example, in the WT SARS-CoV-2 FCD sequence PRRARS, the R at position 5 corresponds to P1, the S to position P1’. We will note the FurinDB identification parenthetically.

We first applied AlphaFold (*11*) through the ColabFold (*13*) environment to examine how well we could match the folded structure of furin. The result is shown in Fig. 1a. The AlphaFold structure for furin matches that from the PDB entry 6EQX (*10*) to within a root mean square deviation of 1.79Å. Next, we included the furin inhibitor RRRVR-4-aminomethyl-benzamidine (RRRVR-Amba) from Ref. (*10*) into the AlphaFold Multimer program (*12*), but because we could not enter the nonstandard residue Amba into the AlphaFold search we substituted tyrosine, which is similar to Amba away from the side chain terminus. As shown in Fig. 1b, this produces a structure substantially similar to furin with bound RRRVR-Amba, except that the S1 pocket, which binds the position 6(P1) Amba nonstandard residue, accepts the position 5(P2) arginine for RRRVRY. In essence, the Y for Amba substitution shifts the sequence to P5-P4-P3-P2-P1-P1’. The RMSD deviation of the RRRVR-Amba backbone from the RRRVRY backbone in the binding position is 2.77Å, which is relatively small and reasonable given the different placement of the Amba vs. arginine in the S1 pocket.

This sequence is very similar to the SARS-CoV-2 delta sequence commencing with arginine at 681, namely, RRRARS. It is known that the furin cleaves between the arginine at 685 and serine at 686. Hence, we hypothesize that the fifth residue(P1) enters the furin S1 pocket. When we utilize AlphaFold Multimer to explore the binding of the WT sequence (beginning at 681) PRRARS, we do find that the fifth arginine(P1) enters the furin S1 pocket, and binds strongly to two aspartic acid residues at positions 258 near the pocket entry, and 306 at the interior end of the pocket (Fig. 1D). We note that the last arginine in the RVKR sequence of Ref. (*8*) also has close proximity to D258 and D306. The RMSD deviation of the RRRVR-Amba backbone from the PRRARS backbone of the FCD for SARS-CoV-2 from AlphaFold Multimer is 4.1Å. This is not surprising given the large sequence difference.

Arginine is particularly well suited for this binding, with its three side chain nitrogens in contrast to the single nitrogen in lysine. A lysine at the fifth position (P1) is only able to bind to the D306. Of all 62 observed sequences identified from GISAID for SARS-CoV-2, 58 have an arginine at position 5 (P1). For the other human coronaviruses, four (MERS, OC42, HUK1a,b) have an arginine at this position. NL63 and 229E have serines at this position, and the H5N1 flu has lysine.

There is also a strong bias towards a hydrophobic residue at position 4(P2) in the SARS-CoV-2 sequences. Alanine arises there in 46 of 62 sequences, and valine in 4 of 62. The alanine side chain carbon is within 5Å of side chain carbons on W147 and L120 from the furin in the delta structure. Of the other viruses, MERS and 229E have valine at position 4, while the others have arginine (OC43, HUK1a,b) or serine (NL63) at this position. The H5N1 flu has lysine at this position.

Given a starting structure, we can simulate and measure characteristics of the binding, such as counting FCD-furin HBonds, calculating the binding energy of the complex, or measuring the interfacial surface area, defined as half the difference between the solvent accessible surface area of the separated furin and FCD vs the solvent accessible surface area of the complex. We utilize the `YASARA` molecular modeling program (*16*), simulating each bound FCD-furin complex for at least 10 ns past energy minimization and equilibration. We then count interfacial protein hydrogen bonds using the criteria outlined for `YASARA` (*17*). For computing the binding free energy, we use the Generalized Born Surface Area (GBSA) endpoint free energy calculation from the `HawkDock` server (*18*). Because the binding interface is tight, there is essentially no water entry between the peptide and furin. As shown in the supplemental information, we obtain a strong correlation between the GBSA binding energy and the FCD-furin HBond count (Fig. S1). For the rest of this paper, we shall use the FCD-furin HBond count as a proxy for binding strength. Note that in this approach, salt bridges of proximate residues, are effectively counted as H-bonds between basic side chain amide groups and acidic side chain carboxyl groups. Hence, the R685 residue of the spike protein FCD forms a salt bridge with the D306 residue of the furin protein, but this is counted in FCD-furin HBonds in this approach.

We have used AlphaFold as the only way to generate a candidate structure for the binding of the WT peptide to furin. With the other sequences we have a choice of using AlphaFold or using the mutation approach within YASARA. We generally find that there are small differences in favor of the mutation approach as we discuss in detail in the supplemental information (Fig. S2).

We have surveyed a total of 62 observed six member SARS-CoV-2 furin FCD sequences at 681-686 for this paper drawn from from the GISAID database (*19*–*21*), out of which the delta sequence RRRARS is the top binder to within statistical significance. Those used in this paper are acknowledged in Supplemental Table 1. Fig. 2A shows the FCD-furin HBond counts for the prevalent 681-686 sequences WT(PRRARS), delta (RRRARS), and omicron/alpha (HRRARS). We have also included KSHRRA as an alternate omicron sequence in view of the N679K mutation. Additionally, we include seven observed but rare sequences found from GISAID chosen either for their frequency of occurrence or their high FCD-furin HBond count. Finally, we include two sequences (PRRDRS and KRRARS) which can be arrived at by two base mutations from either the WT or delta variants. The prevalent codon at position 684 cannot swap by a single base to obtain D, and the prevalent codon at position 681 cannot swap by a single base to obtain K. By performing pairwise t-tests within GraphPad, we find that the FCD-furin HBond count for the delta variant sequence binding to furin exceeds all but one of the observed sequences with statistical significance (*p* < 0.05) or extreme (*p* < 0.0001) statistical significance, and differences are statistically insignificant in comparison to the observed SRRARS and unobserved PRRDRS and KRRARS sequences (p > .1 for each).

A similar picture emerges compared to other human coronaviruses and the H5N1 flu as shown in Fig. 2b. The candidate sequences for OC43, NL63, HUK1a,b, 229E, and MERS were obtained by homology alignment of the spike proteins using BLAST (*22*). The FCD-furin HBond count difference between the delta variant and these viral sequences is extremely significant (p < 0.0001). We note that the binding is strongest for the cold viruses HUK1a,b and OC43.

To assess the importance of the different residues in the six member peptide to binding strength, we analyzed the hydrogen bonding patterns in detail. We display a heat map in Fig. 3 for many of the sequences shown in Fig. 2. We find in nearly every case that the strongest binding, representing a significant plurality of the binding strength, is for the position 5(P1) residue, with arginine the preferred amino acid there. Notably, the H5N1 sequence with a K at position 5, and the trial sequence PKKAKS where all arginines are replaced by lysines, fare poorly at position 5(P1) compared to the other sequences.

In searching for an understanding for these observations, we have uncovered three correlations, two of which that can independently explain nearly 50% of the variation between SARS-CoV-2 sequences and separately between viruses. First, by examining the root mean square fluctuation of the backbone C-alpha of the first residue (P5), compared to the FCD-furin HBond counts of the observed sequences with at least 50 appearances in the GISAID tables for SARS-CoV-2, we see in Fig. 4A that this backbone fluctuation correlates inversely with the binding strength with a linear regression coefficient of *R*^2^ = 0.53. Fig. 4B shows the correlation between the FCD-furin HBond count and the RMSF of the first alpha carbon (CA) atom for delta, the six other human coronaviruses with homology in this region, and the H5N1 flu virus. The linear regression coefficient is *r*^2^ = 0.49. The best fit slope of −2.74±1.25 FCD-furin HBonds/Å is less than that for SARS-CoV-2 (−4.33± 1.54 FCD-furin Hbonds/Å), but the difference is statistically insignificant. Second, by examining the number of FCD-furin HBonds associated with the residue at position 5(P1), we observe (Figs. 4C, 4D) that there is a high degree of correlation with the total FCD-furin HBond count. For the observed higher frequency furin binding sequences, the best fit slope is 1.41±.38 with *R*^2^ = 0.59, and for comparison of delta to other viruses, the slope is similar 1.77±.51 with *R*^2^ = 0.66. Third, as shown in Fig. 4E, the number of basic residues (H,K, or R) in the six residue sequence helps determine the maximum number of FCD-furin HBonds. Fitting the maximum envelope of the plot to a quadratic, as in Fig. 4F, gives

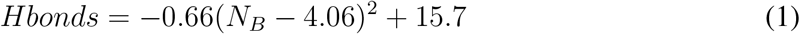

where *N_B_* is the number of bases. The nonlinear regression coefficient is *R*^2^=0.98. This suggests that the maximum number of FCD-furin HBonds is 16, for four basic residues (as per the delta variant), and to within statistical accuracy, no sequence exceeds delta in the number of FCD-furin HBonds to furin.

**Fig. 4.**
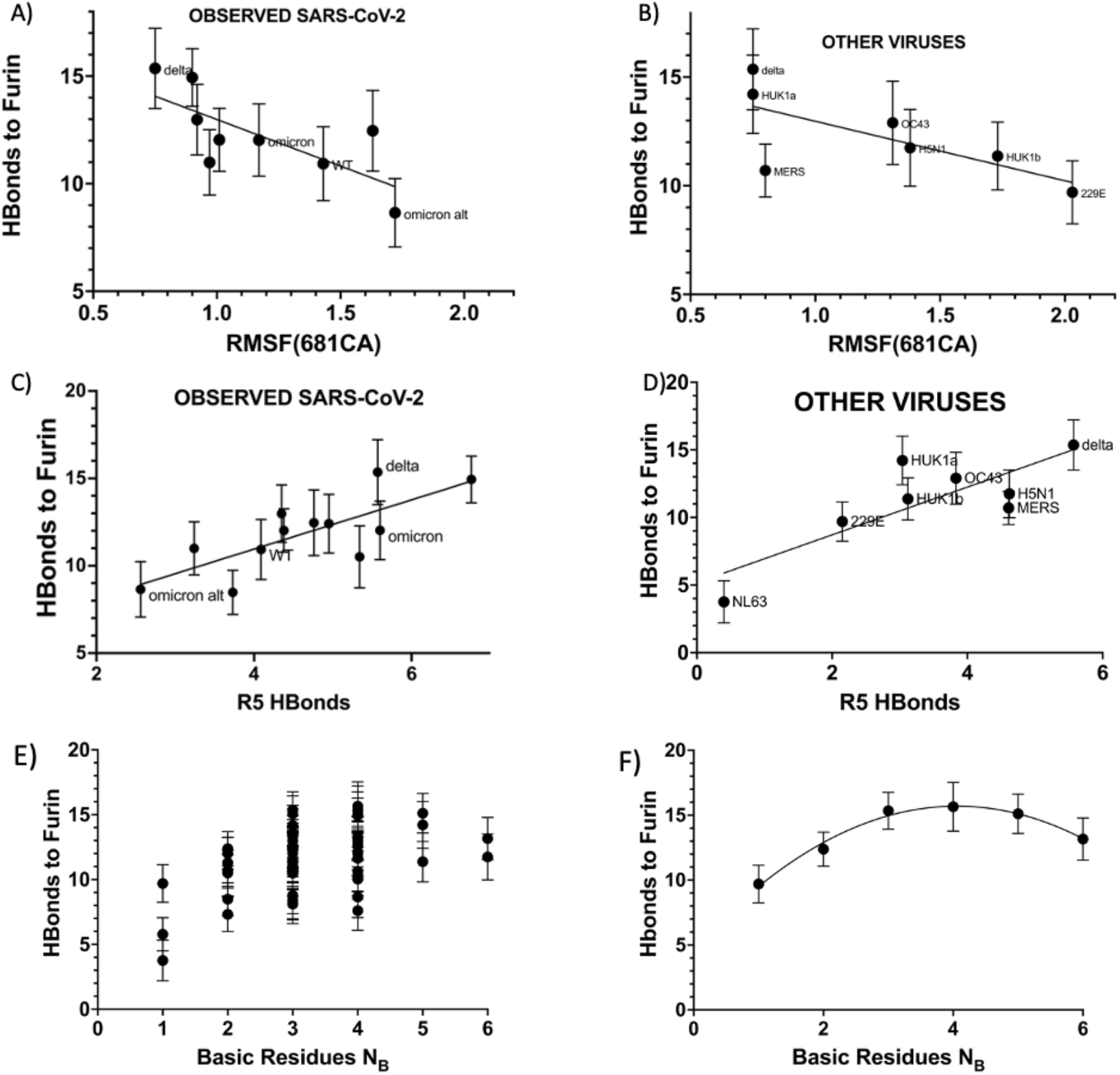
A) Correlation of the backbone fluctuation from Residue 1 of the sequence with the total number of FCD-furin HBonds between the binding sequence and furin for SARS-CoV-2 sequences observed at least 50 times. B) Correlation of the residue 1 backbone fluctuation for delta and other viruses. C) Correlation of the interfacial HBonds for Residue 5 with the total number of HBonds for observed SARS-CoV-2 sequences of A). D) Correlation of the FCD-furin HBonds for Residue 5 with total number of FCD-furin HBonds for delta and other viruses. E) The number of HBonds for a given number of basic FCD residues plotted for 56 sequences. F) The maximum FCD-furin HBonds envelope as a function of the number of basic residues. This is fit with *R*^2^=0.98 by Eq. (1) of the main text. The sequences for the peak values are for 1-6 respectively: PRNSVY (229E coronavirus), PRQARS (SARS-CoV-2), SRRARS (SARS-CoV-2), KRRARS (SARS-CoV-2, unobserved), RRRRRD (Epstein-Barr, ref. (*26*)), RRRRRR (unobserved).

## Discussion

The most important results of this paper are: 1) by using AlphaFold Multimer (*12*) we have validated by comparison to the binding of furin with a known six residue inhibitor, we are able to predict bound structures for over 60 observed FCD sequences of SARS-CoV-2 (at residues 681-686 of the spike protein and two alternate sequences for omicron) and eight other viruses (six human coronaviruses (OC43, HUK1a, HUK1b, MERS, NL63, 299E), the H5N1 influenza, and Epstein-Barr virus). From among these, the delta variant FCD of SARS-CoV-2 has the strongest binding to furin within statistical accuracy, with 15.3 mean FCD-furin hydrogen bonds. 2) Within these sequences we find selection for arginine at position 5 (P1), which fits into a furin S1 pocket having aspartic acids at the entrance and within. The structure of arginine allows binding to both aspartic acids, while lysine’s structure does not. 3) There is also bias towards a hydrophobic residue at position 4(P2) of the six residue FCD, which appears to interface favorably with W147 and L120 of the furin. 4) We find that two features of the sequences each predict about half of the binding strength: (i) the backbone fluctuation of the first residue in the binding sequence correlates inversely with the overall binding strength as measured by FCD-furin HBonds, and (ii) the number of hydrogen bonds associated with the binding of residue 5(P1) in the furin S1 pocket correlates positively with FCD-furin HBond count. This residue never accounts for more than a plurality of the FCD-furin HBonds so it is somewhat surprising that it correlates with the observed trend of binding. (iii) By considering the variation of the FCD sequence with the number of basic residues, we conclude that no more than 16 FCD-furin HBonds are possible, and within statistical accuracy delta achieves the maximum value. We conjecture that the physical basis for this is a tradeoff between binding efficacy of the basic residues (especially arginine) and Coulomb repulsion as more are added.

In conclusion, we find that spike FCD-furin binding depends critically upon insertion of arginine in the fifth position (P1) of the FCD in a furin pocket that includes D258 at the opening and D306 at the interior end. This prediction emerges uniquely from the application of AlphaFold Multimer (*12*) to predict the bound structure, and contrasts with earlier work that employed a docking program for interface prediction (*14*). It is therefore critical to have experimental structural biology test of this prediction.

Note that the omicron FCD sequence is the same as alpha, and alternate FCD sequences (KSHRRA, beginning at K679, or QTQTKS, with K679 at position 5(P1)) have fewer FCD-furin HBonds than any observed variants, consistent with the observed milder impact of omicron on the lungs (*23*–*25*).

We conclude that it is quantitatively unlikely that any SARS-CoV-2 variant, or any other virus can bind significantly more strongly to the furin protease than the delta variant. This is based on a survey of a large number of observed SARS-CoV-2 spike sequences new SARS-CoV-2 spike sequences not yet observed, other human coronaviruses, H5N1 influenza, and Epstein-Barr virus. The basic model for viral infection is that after spike RBD binding to ACE2, furin cleavage at the FCD regulates fusogenicity leading to syncytia and viral reproduction. Our theoretical studies indicate that furin-FCD driven fusogenicity is at its worst with the delta variant among all observed SARS-CoV-2 variants of interest or concern. Of concern and cause for caution are some rarely observed or unobserved FCD sequences which could be just as consequential for furin cleavage as delta (observed: SRRARS, RRRARN,HRRVRS; unobserved: PRRDRS, KRRARS).

## Supporting information

Supplemental information for "Computational Study of the furin cleavage domain..."

## Acknowledgments

We acknowledge fruitful discussions with Victor Muñoz. J.A., M. V, G.G. were partially supported by NSF grant DMS-2030491, G.G. was supported by a grant from the UC Davis Center for Data Science and Artificial Intelligence and by a donation from Protein Architects to J.A and M.V., S. J. was supported by a Md. Abdur Razzaq Global HealthShare Undergraduate Research Fellowship.

## Supplementary materials

Materials and Methods

Supplementary Text

Figs. S1, S2, S3

Tables ST1

## References

1. E. C. Holmes, et al., Cell 184, 4848 (2021).

2. T. P. Peacock, et al., bioRxiv p. 2020.09.30.318311 (2020).

3. T. P. Peacock, et al., Nature Microbiology 6, 899 (2021).

4. A. Saito, et al., Nature (2021).

5. Y. Wu, S. Zhao, Stem Cell Research 50, 102115 (2021).

6. P. Decha, et al., Biophysical Journal 95, 128 (2008).

7. D. J. Benton, et al., Nature 588, 327 (2020).

8. S. Henrich, et al., Nature Structural & Molecular Biology 10, 520 (2003).

9. M. M. Kacprzak, et al., Journal of Biological Chemistry 279, 36788 (2004).

10. S. O. Dahms, K. Hardes, T. Steinmetzer, M. E. Than, Biochemistry 57, 925 (2018).

11. J. Jumper, R. Evans, A. e. a. Pritzel, Nature (2021).

12. R. Evans, et al., bioRxiv p. 2021.10.04.463034 (2021).

13. M. Mirdita, S. Ovchinnikov, M. Steinegger, bioRxiv p. 2021.08.15.456425 (2021).

14. N. Vankadari, Journal of Physical Chemistry Letters 11, 6655 (2020).

15. S. Tian, Q. Huang, Y. Fang, J. Wu, International Journal of Molecular Sciences 12 (2011).

16. E. Krieger, G. Vriend, Bioinformatics 30, 2981 (2014).

17. E. Krieger, R. L. Dunbrack, R. W. Hooft, B. Krieger, Computational Drug Discovery and Design (Springer, 2012), pp. 405–421.

18. G. Q. Weng, et al., Nucleic Acids Research 47, W322 (2019).

19. S. Khare, et al., China CDC Weekly 3, 1049 (2021).

20. S. Elbe, G. Buckland-Merrett, Global Challenges 1, 33 (2017).

21. Y. Shu, J. McCauley, Eurosurveillance 22, 30494 (2017).

22. E. W. Sayers, et al., Nucleic Acids Research p. gkab1112 (2021).

23. B. Meng, et al., bioRxiv p. 2021.12.17.473248 (2021).

24. Z. Cong, et al., bioRxiv p. 2021.12.16.472934 (2021).

25. M. Z. Jawaid, A. Baidya, R. Mahboubi-Ardakani, R. L. Davis, D. L. Cox, bioRxiv p. 2021.12.14.472704 (2021).

26. E. Braun, D. Sauter, Clinical & Translational Immunology 8, e1073 (2019).

