## Supplemental information for "Computational Study of the furin cleavage domain..." for "Computational study of the furin cleavage domain of SARS-CoV-2: delta binds strongest of extant variants"

M. Zaki Jawaid,[1] A. Baidya,[1] S. Jakovcevic,[1,3]
J. Lusk[2], R. Mahboubi-Ardakani,[1], N. Solomon,[2] G. Gonzalez,[2]
J. Arsuaga,[2,3] M. Vazquez,[2,4] R.L. Davis,[5], D.L. Cox,[1,5*]
[1]Department of Physics, University of California, Davis, USA
[2] Department of Mathematics, University of California, Davis, USA
[3]Department of Cellular and Molecular Biology, University of California, Davis, USA
[4]Department of Microbiology and Molecular Genetics, University of California, Davis, USA
[5]Protein Architects Corp., Penn Valley, PA, USA

### Materials and Methods

#### 0.1 Molecular Models

For the furin structure and for furin binding to the inhibitor RRRVR-Amba, we used PDB entry 6EQX (*1*).

#### 0.2 Sequence Alignment for Other Coronaviruses

To identify homology in the furin cleavage domain for the other coronaviruses OC43, NL63, 229E, MERS, HUK1a, HUK1b, we utilized BLAST (*2*) at the National Center for Biotechnology Information. We compared entire spike sequences and zoomed in on the furin cleavage domain based upon the PRRARS sequence for SARS-CoV-2.

#### 0.3 Identification of other SARS-CoV-2 sequences using GISAID

#### 0.4 Genomic Data Set and Sequence Pre-Processing

We obtained SARS-CoV-2 sequences for this study from the GISAID database on Nov 11, 2020 (*3*). Our data set contains FASTA files for every complete human SARS-CoV-2 nucleotide sequence (from all geographical locations) available in GISAID between and 12/1/19 and 7/11/2021. The sequences were then aligned using `ClustalOmega` with the default parameters (*4*). We found that `ClustalOmega` ran faster on our data set than common alternatives like `ClustalW` (*5*) and `MUSCLE` (*6*).

After aligning the sequences, we extracted the spike protein by comparing the aligned sequence with the NCBI's SARS-CoV-2 reference sequence (NC_045512.2; "WT") (*7*) and tabulated the frequencies of different furin binding domain inserts.

Table ST1 includes accession numbers and acknowledgements for the first of each 111 unique nucleotide sequences referenced in this paper as they appear in GISAID.

#### 0.5 Molecular Dynamics

To simulate the protein-protein interactions, we used the molecular-modelling package `YASARA` (*8*) to substitute individual residues and to search for minimum-energy conformations on the resulting modified structures of the FCD-furin. For all of the structures, we carried out an energy-minimization (EM) routine, which includes steepest descent and simulated annealing (until free energy stabilizes to within 50 J/mol) minimization to remove clashes. All molecular-dynamics simulations were run using the AMBER14 force field with (*9*) for solute, GAFF2 (*10*) and AM1BCC (*11*) for ligands, and TIP3P for water. The cutoff was 8 Å for Van der Waals forces (AMBER's default value (*12*)) and no cutoff was applied for electrostatic forces (using the Particle Mesh Ewald algorithm (*13*)). The equations of motion were integrated with a multiple timestep of 1.25 fs for bonded interactions and 2.5 fs for non-bonded interactions at $T = 298$ K and $P = 1$ atm (NPT ensemble) via algorithms described in (*14*). Prior to counting the FCD-furin hydrogen bonds and calculating the free energy, we carry out several pre-processing steps on the structure including an optimization of the hydrogen-bonding network (*15*) to increase the solute stability and a $pK_{\mathrm{a}}$ prediction to fine-tune the protonation states of protein residues at the chosen pH of 7.4 (*14*). Insertions and mutations were carried out using YASARA's BuildLoop and SwapRes commands (*14*) respectively. Simulation data was collected every 100ps after 1-2ns of equilibration time, as determined by the solute root mean square deviations (RMSDs) from the starting structure. For all bound structures, we ran for at least 10 ns post equilibrium, and verified stability of time series for FCD-furin hydrogen bond counts and root mean square deviation (RMSD) from these starting structure. Because of concerns about the validity of short time simulations, and more variability for the weaker binding for the omicron RBD-ACE2 complex, we ran for 40 ns postequilibration in that case.

The FCD-furin hydrogen bond (HBond) counts were tabulated using a distance and angle approximation between donor and acceptor atoms as described in (*15*). Note that in this approach, salt bridges of proximate residues, are effectively counted as H-bonds between basic side chain amide groups and acidic side chain carboxyl groups. Hence, the R685 residue of the spike protein FCD forms a salt bridge with the D306 residue of the furin protein, but this is counted in HBonds in this approach.

Note that in view of the likely ambient pH for cell surface or endosomal furin cleavage, and the polybasic environment of the FCD, we have assumed all histidines to be singly protonated at the delta site. Choosing the epsilon site makes little difference. For the alpha sequence, doubly protonated histidine binds more strongly, but for the alternate omicron sequence, there is little difference among the three protonation states.

#### 0.6 Endpoint Free Energy Analysis

We calculated binding free energy for the energy-minimized structure using the molecular mechanics/generalized Born surface area (MM/GBSA) method (*16–18*), which is implemented by the `HawkDock` server (*19*). While the MM/GBSA approximations overestimate the magnitude of binding free energy relative to *in-vitro* methods, the obtained values correlate well with H-bond counts. For each RBD-ACE2, RBD-AB, and NTD-Ab binding pair we average over five snapshots of equilibrium conformations. For each FCD-furin pair, we average over ten snapshots of equilibrium conformations.

#### 0.7 Use of ColabFold/AlphaFold for Furin binding domain

Full details of this method are provided in (*20–22*). In brief, we used the heterocomplex prediction method known as AlphaFold-Multimer (*20, 21*) as implemented within ColabFold (*22*) to predict the best bound structure to the furin enzyme of the six residue FCD from the WT protein. We inferred the ordering of this sequence by comparison with a very similar six residue peptide inhibitor of furin with the sequence RRRVR-aminomethyl-benzamidine (RRRVR-Amba) (*1*). In this case the backbone of the WT FCD aligns well with that of the inhibitor, but the arginine at residue 5 enters the furin S1 pocket (*1*) while the Amba enters the furin S1 pocket for the inhibitor. The serine is in proper cleavage position for furin. Most other structures were then obtained by mutation from the predicted WT FCD-furin structure.

#### 0.8 Statistical Analysis and Graphics

We computed the statistical significance of pairwise differences using the GraphPad unpaired t-test calculator. Regression analysis for Fig. 4, Fig. S1 was carried out using the GraphPad Prism (v. 9) package. Structural images for Fig. 1 were created in YASARA. Figs. 2,4, S1, S2 were created with GraphPad Prism (v. 9). Fig. 2 was created with Seaborn (v. 0.11.2), a Python data visualization library.

### Supplementary Text

#### 0.9 Correlation of FCD-furin HBonds and MM/GBSA binding energy estimates

Fig. S1 summarizes the correlation between FCD-furin HBonds and the MM/GBSA estimates for binding energy in kcal/mole. Note that MM/GBSA usually overestimates binding energy strength significantly but is good for producing binding energy trends. The regression coefficient is $R^2 = 0.61$, and the best fit slope is -.107 Hbonds/(Kcal/mole) with 95$\%$ confidence intervals of -.110 to -.1042. Clearly, the correlation is strong between FCD-furin HBonds and binding energy.

#### 0.10 Differences between simulations with AlphaFold and mutation

While we must take the WT FCD-furin structure from AlphaFold (*20, 21*), we can mutate using the Swap command in YASARA from there to obtain other starting structures for molecular dynamics simulations. In general, AlphaFold produces structures with slightly less binding strength than mutating from the WT, with a few exceptions, the delta variant being one. This is demonstrated for five sequences in Fig. S2. Accordingly, because the resultant binding is stronger we have used the mutant results where possible to provide a more accurate starting point for the equilibration runs in molecular dynamics.

#### 0.11 Examples of sequence frequency and codons

Fig. S3 shows a table of FCD sequences used in the figures as well as one synonymous/silent mutation based upon the consensus codons for WT, alpha, and delta. The last two entries are for unobserved but potentially potent FCD sequences. Mutation to those would require two base swaps from either WT or delta.

### References


1. S. O. Dahms, K. Hardes, T. Steinmetzer, M. E. Than, *Biochemistry* **57**, 925 (2018).
2. E. W. Sayers, *et al.*, *Nucleic Acids Research* p. gkab1112 (2021).
3. Y. Shu, J. McCauley, *EuroSurveillance* **22** (2017).
4. F. Sievers, D. G. Higgins, *Protein Science* **27**, 135 (2018).
5. J. D. Thompson, D. G. Higgins, T. J. Gibson, *Nucleic Acids Research* **22**, 4673 (1994).
6. R. C. Edgar, *Nucleic Acids Research* **32**, 1792 (2004).
7. F. Wu, *et al.*, *Nature* **579**, 265 (2020).
8. E. Krieger, G. Vriend, *Bioinformatics* **30**, 2981 (2014).
9. J. A. Maier, *et al.*, *Journal of chemical theory and computation* **11**, 3696 (2015).
10. J. Wang, R. M. Wolf, J. W. Caldwell, P. A. Kollman, D. A. Case, *Journal of computational chemistry* **25**, 1157 (2004).
11. A. Jakalian, D. B. Jack, C. I. Bayly, *Journal of computational chemistry* **23**, 1623 (2002).
12. V. Hornak, *et al.*, *Proteins: Structure, Function, and Bioinformatics* **65**, 712 (2006).
13. U. Essmann, *et al.*, *The Journal of chemical physics* **103**, 8577 (1995).
14. E. Krieger, G. Vriend, *Journal of computational chemistry* **36**, 996 (2015).
15. E. Krieger, R. L. Dunbrack, R. W. Hooft, B. Krieger, *Computational Drug Discovery and Design* (Springer, 2012), pp. 405–421.

16. T. Hou, J. Wang, Y. Li, W. Wang, *Journal of Chemical Information & Modeling* **51**, 69 (2001).

17. H. Sun, Y. Li, S. Tian, L. Xu, T. Hou, *Physical Chemistry Chemical Physics* **16**, 16719 (2014).

18. F. Chen, *et al.*, *Physical Chemistry Chemical Physics* **18**, 22129 (2016).

19. G. Q. Weng, *et al.*, *Nucleic Acids Research* **47**, W322 (2019).

20. J. Jumper, R. Evans, A. e. a. Pritzel, *Nature* (2021).

21. R. Evans, *et al.*, *bioRxiv* p. 2021.10.04.463034 (2021).

22. M. Mirdita, S. Ovchinnikov, M. Steinegger, *bioRxiv* p. 2021.08.15.456425 (2021).

#### Table ST1: GISAID accession numbers and acknowledgements

We have included the acknowledededgements and access numbers for the first appearance in base notation of the sequences referenced in this paper (111).

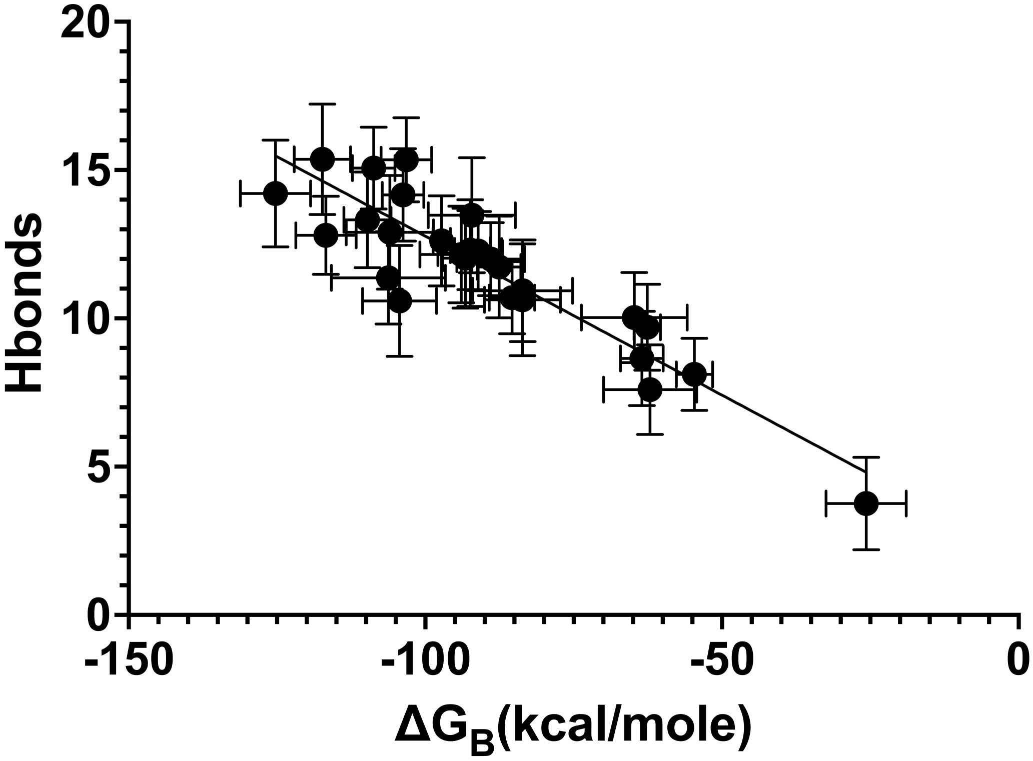


Figure 1: **Correlation of FCD-furin HBond counts with MM/GBSA Binding Energy** FCD-furin HBond counts are estimates from YASARA (*8,15*) simulations, while MM/GBSA Binding Energy comes from the HawkDock server (*19*). Regression analysis using GraphPad Prism 9 provides the straight line fit (see text for details).

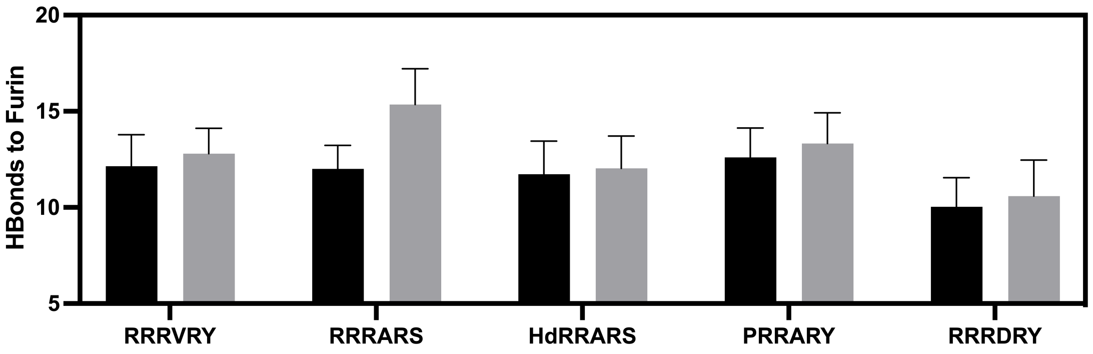


Figure 2: **Differences in equilibrated FCD-furin HBond counts between AlphaFold generated starting FCD-furin structures and starting structures mutated from the AlphaFold WT structure** In general, after equilibration the AlphaFold structures have slightly less binding strength, with a few exceptions such as the delta variant where AlphaFold misses dramatically. For comparison, the $p$-values for AlphaFold vs mutant in this plot are RRRVRY- $p$=0.0035 (very significant); RRRARS (delta)- $p$¡0.000001 (extremely signficant); HRRARS (alpha/omicron)- $p$=0.181 (not significant); PRRARY - $p$=0.00094 (very significant); RRRDRY - $p$=0.0164 (very significant)

| Frequency | P1 | P2 | P3 | P4 | P5 | P6 |
|---|---|---|---|---|---|---|
| 47.40000000% | H(CAT) | R(CGG) | R(CGG) | A(GCA) | R(CGT) | S(AGT) |
| 5.90000000% | R(CGT) | R(CGG) | R(CGG) | A(GCA) | R(CGT) | S(AGT) |
| 45.60000000% | P(CCT) | R(CGG) | R(CGG) | A(GCA) | R(CGT) | S(AGT) |
| 0.07247210% | H(CAT) | R(CGG) | R(CGT) | A(GCA) | R(CGT) | S(AGT) |
| 0.04050000% | H(CAT) | R(CGG) | R(CGG) | V(GTA) | R(CGT) | S(AGT) |
| 0.02800000% | P(CCT) | R(CGG) | R(CGG) | V(GTA) | R(CGT) | S(AGT) |
| 0.02710000% | L(CTT) | R(CGG) | R(CGG) | A(GCA) | R(CGT) | S(AGT) |
| 0.00425500% | Y(TAT) | R(CGG) | R(CGG) | A(GCA) | R(CGT) | S(AGT) |
| 0.00397982% | S(TCT) | R(CGG) | R(CGG) | A(GCA) | R(CGT) | S(AGT) |
| 0.00001474% | P(CCT) | R(CGG) | R(CGG) | E(GAA) | R(CGT) | S(AGT) |
| 0.00000000% | P(CCT) | R(CGG) | R(CGG) | D(GAC) | R(CGT) | S(AGT) |
| 0.00000000% | K(AAA) | R(CGG) | R(CGG) | A(GCA) | R(CGT) | S(AGT) |

Figure 3: **Examples of FCD sequences from GISAID for analysis here with codons** The observed frequencies of sequences between 12/1/19 and 7/11/21 appear at left, and the predominant codons for each position are tabulated. Row 4 shows a synonymous/silent mutation to the alpha variant, while the rest show missense mutations. The last two sequences are unobserved (requiring double codon swaps relative to either WT or delta) but bind as strongly to furin as the delta FCD. Note that over this entire pre-omicron time frame that delta (RRRARS) has less accumulated percentage of the sequences than WT (PRRARS) or alpha (HRRARS).
